# Race modifies default mode network connectivity in Alzheimer’s disease

**DOI:** 10.1101/472910

**Authors:** Maria B. Misiura, Junjie Wu, Deqiang Qiu, Jennifer C. Howell, Monica W. Parker, Jessica A. Turner, William T. Hu

## Abstract

**Objective:** To determine if resting state functional MRI biomarkers of Alzheimer’s disease (AD) differ between older African Americans and Caucasians.

**Methods:** We analyzed MRI profiles from 78 individuals (31 African Americans, 47 Caucasians) with normal cognition (n=36) or mild cognitive impairment/mild AD dementia (MCI/AD, n=42). We compared AD-associated intra-network functional connectivity of the default mode network (DMN) according to race, and correlated domain-specific cognitive functions with functional connectivity which differed between the racial groups.

**Results:** We identified key differences in DMN functional connectivity associated with AD between the races. Whereas MCI/AD was associated with decreased functional connectivity within the midline core DMN subsystem in older Caucasians, MCI/AD was instead associated with increased functional connectivity within the same subsystem of older African Americans. This is despite decreased functional connectivity in the medial temporal lobe DMN subsystem in both races. Memory function was also positively associated with connectivity between the precuneus and the posterior cingulate/inferior parietal lobule within the midline core subsystem, in keeping with a less amnestic-profile in older African Americans with MCI/AD.

**Conclusions:** These findings provide structural support that race modifies the AD phenotypes downstream from cerebral amyloid deposition, and suggests a rsf-MRI correlate of African American’s less amnestic neuropsychological profile in MCI/AD.

## Introduction

Older African Americans are twice as likely to develop Alzheimer’s disease (AD) as older non-Hispanic Caucasians.^1^ Epidemiological and genome-wide association studies in clinically defined cases of dementia suggest that AD may manifest differently according to race, vascular risks, educational duration, and susceptibility genes.^1,2^ We previously reported that AD was associated with lower CSF tau-related biomarker levels in African Americans than Caucasians, despite similar levels of beta-amyloid 1-42 (Aβ42).^2^ We did not find a difference in hippocampal volumes, but it is not known if other objective biomarkers differed between the two races. Here we analyze MRI biomarkers that may reflect earlier AD-related processes and may better inform the relationship between race and AD neuroimaging phenotypes.

In AD, disrupted functional connectivity in the default mode network (DMN) is reported to occur following amyloid biomarker changes but before significant hippocampal atrophy.^3,4^ We hypothesized that, similar to baseline and AD-associated differences in CSF tau biomarkers between the two racial groups, older African Americans also had AD-associated DMN alterations distinct from older Caucasians. To test this hypothesis, we analyzed resting state functional (rsf-) MRI from the same group of subjects with normal cognition (NC) or early AD (mild cognitive impairment or mild AD dementia, MCI/AD), and analyzed if race, diagnosis, and other factors influenced DMN connectivity measures.

## Methods

### Standard Protocol Approvals, Registrations, and Patient Consents

This study was approved by the Emory University Institutional Review Board and written consent was obtained from all subjects.

### Participants

We included 78 participants who underwent uniform clinical, neuropsychological, genetic, and CSF biomarker analysis as previously described (Table e-1).^2^ MRI analysis was performed using a modified Alzheimer’s Disease Neuroimaging Initiative (ADNI) protocol (e-Methods). For functional connectivity, regions of the DMN include the precuneus (PCUN), posterior cingulate gyrus/inferior parietal lobule (PCC/IPL), the ventromedial prefrontal cortex (vmPFC), and the hippocampus. We used a data driven approach (Independent Component Analysis; ICA) using the Group ICA of fMRI Toolbox (GIFT) to identify large-scale brain networks, and calculated connectivity measures between DMN nodes.^5^

**Table 1.** Effects of race, MCI/AD, and risk genes/factors on DMN connectivity. *Note: N.S. = not significant*

|  | vmPFC-PCUN<br>(F, p) | vmPFC-PCC/IPL<br>(F, p) | PCUN-PCC/IPL<br>(F, p) |
| --- | --- | --- | --- |
| Race x MCI/AD | 3.150 (0.02) | 3.091 (0.01) | 13.700 (<0.001) |
| *APOE* ε4 | 4.079 (0.047) | 3.499 (0.065) | N.S. |
| *ABCA7* | 3.511 (0.065) | N.S. | 5.328 (0.024) |
| Hypertension | 3.063 (0.084) | N.S. | N.S. |
| Diabetes | N.S. | N.S. | N.S. |
| Diabetes x MCI/AD | N.S. | 2.497 (0.067) | 3.009 (0.056) |

### Data availability

The data that support the findings of this study are available from the corresponding author, upon reasonable request.

### Statistical Analyses

Statistical analysis was performed in R version 3.3.3. We constructed multivariate multiple linear regression models with intra-network connectivity between the three posterior DMN nodes as the dependent variables; race, sex, and diagnosis (normal cognition, MCI/AD), age, and mean framewise displacement (MFWD) as independent variables, as well as a higher order interaction term (race X diagnosis). Measures of connectivity between the anterior DMN component (vmPFC) and the hippocampus, PCUN, and PCC/IPL were similarly analyzed for race-associated differences. We additionally examined the relationship of potential confounding effects of *APOE* ε4, *ABCA7* risk allele, hypertension, and diabetes which differed between the races.^2^

To determine possible functional consequences of race-associated differences in DMN connectivity, we also correlated DMN connectivity that differed between the races with domain-specific cognitive functions. In this model, domain-specific Z-scores for executive, memory, language, and visual spatial function were the dependent variable, and diagnosis (NC vs. MCI/AD), connectivity, age, and having at least one *APOE* ε4 allele were the independent variables.

## Results

### Race modifies the relationship between MCI/AD and midline core DMN connectivity

Consistent with published results, MCI/AD was associated with reduced anterior-posterior DMN connectivity (vmPFC-PCC/IPL and vmPFC-PCUN) and unchanged PCUN-PCC/IPL connectivity in older Caucasians, adjusting for age and gender (Fig 1A). Race modified the relationship between MCI/AD diagnosis and connectivity for all three examined midline node pairs. Unlike in Caucasians, MCI/AD was associated with *increased* PCUN-PCC/IPL connectivity in African Americans, and unchanged anterior-posterior DMN connectivity. Further adjustment for *ABCA7* and *APOE* genotypes showed similar results (Table 1).

**Figure 1.**
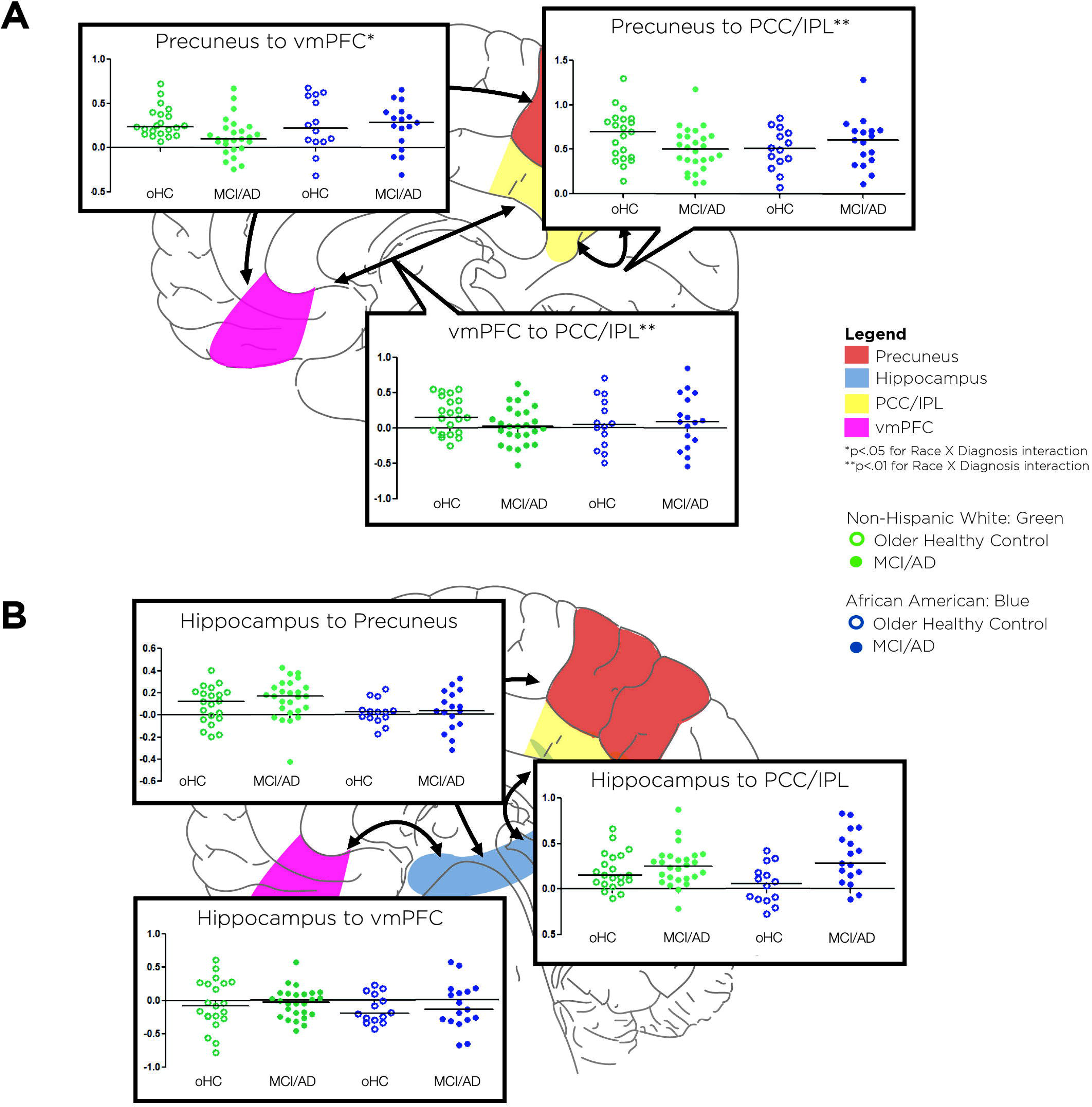
Connectivity and diagnosis. Y-axes for all graphs are functional connectivity. Bars represent medians for each group. Panel A depicts connectivity and diagnosis relationships among the PCUN, PCC/IPL, and vmPFC nodes. MCI/AD was associated with reduced anterior-posterior DMN connectivity (vmPFC-PCC/IPL and vmPFC-PCUN) in Caucasians but not African Americans. In contrast, MCI/AD was associated with *increased* PCUN-PCC/IPL connectivity in African Americans but not Caucasians. Panel B depicts connectivity between the hippocampi and midline core DMN nodes. Unlike in A, MCI/AD was associated with reduced connectivity between the hippocampus and midline core nodes irrespective of race.

### Race does not modify the relationship between MCI/AD and medial temporal DMN connectivity

Also consistent with published results, MCI/AD was associated with reduced connectivity involving the hippocampus in Caucasians (adjusting for age and gender, Fig 1B; hippocampus-PCUN, *F (2*, 75)=2.13, *p*=0.05; hippocampus-PCC/IPL, *F*(2, 75)=2.17, *p*=0.04). Race did not influence hippocampal DMN connectivity, nor did it modify the relationship between MCI/AD and hippocampal connectivity.

### Greater vmPFC-PCUN connectivity is associated with better memory

Finally, we determined if the connectivity profile differences between African Americans and Caucasians were associated with distinct neuropsychological profiles. Across the entire cohort, greater PCUN-PCC/IPL connectivity correlated with better memory functions independent of race (Table 2). Because increased PCUN-PCC/IPL connectivity was associated with MCI/AD only in African Americans, these results together are in keeping with less amnestic MCI/AD profiles (at the group level) in African Americans than Caucasians.

**Table 2.** Effects of diagnosis, risk genes/factors, and DMN connectivity on memory.

| Variable | Coefficient (95% confidence interval) | P |
| --- | --- | --- |
| MCI/AD diagnosis | -1.362 (-1.942, -0.782) | <0.001 |
| Having at least one *APOE* ε4 allele | -0.861 (-1.441, -0.280) | 0.004 |
| Age | 0.066 (0.026, 0.105) | 0.001 |
| PCUN-PCC/IPL connectivity | 1.235 (0.124, 2.347) | 0.030 |

## Discussion

The biological underpinning for epidemiologic and phenotypic differences in AD between older African Americans and Caucasians remains poorly defined. Here we show that race altered MCI/AD-associated connectivity profiles in the midline core DMN subsystem but not the amyloid-associated medial temporal subsystem. These differences cannot be readily explained by genetic risks from *ABCA7* and *APOE* alone, but likely have clinically meaningful impact as increased posterior midline DMN connectivity in African Americans with MCI/AD corresponds to their less amnestic neuropsychological profiles.

Race-related AD phenotypic differences have historically been attributed to education quality, access to health care, and cerebrovascular disease.^6^ The lower CSF tau levels in older African Americans regardless of cognitive status cannot be adequately explained by these explanations,^2^ and may correlate with midline DMN connectivity profiles. It remains controversial whether altered midline core connectivity reflects post mortem or *in vivo* measures of tau pathology,^8^ but this subsystem correlates only marginally with cerebral amyloid deposition.^7^ Increased DMN connectivity was previously linked to non-AD neurodegenerative diseases, and it is possible that greater non-AD co-pathology in African American AD cases on autopsy begins early rather than late following cerebral amyloid deposition.^9^ Alternatively, increased connectivity may represent network compensation, which suggests greater – not less – cognitive reserve in African Americans with AD to account for their slower clinical progression.^10^ Because there is significant overlap between the groupings, future work should prospectively analyze the relationship between tau PET tracer retention, CSF tau levels, and DMN connectivity in both Caucasians and African Americans. Because African Americans with MCI/AD also have a slower cognitive decline than Caucasians, PCUN-PCC/IPL connectivity – increased at the group level in African Americans – should also be tested as a prognostic biomarker in longitudinal studies.

While these findings extend our previous description of a specific AD endophenotype in African Americans, this study was limited by the number of rsf-MRI scans of insufficient quality, the association nature of the relationships between race and connectivity, and lack of neuropathologic correlate for connectivity differences. Nevertheless, when interpreted with previous data, this study lends further support to the notion that amyloid-related changes –CSF Aβ42 level, and medial temporal DMN connectivity – are common to African Americans and Caucasians in AD, but tau-related changes – CSF tau levels, midline core DMN connectivity, and clinical progression – are modified by race. Not only do these findings promote caution when generalizing AD diagnostic biomarkers developed in Caucasians to African Americans, they may also help refine the subject selection process for future disease modifying therapies.

## Acknowledgements

The authors acknowledge the following persons for assistance with enrollment: James Lah, MD, PhD; Allan Levey, MD, PhD; Chadwick Hales, MD, PhD; Angela Ashley, MD; Felicia Goldstein, PhD; and Andrea Kippels, CNP.

## Appendix 1. Author Contributions

| Name | Location | Role | Contribution |
| --- | --- | --- | --- |
| Maria B. Misiura, M.A. | Psychology Department, Georgia State University. Atlanta, GA | Author | Designed and conceptualized study; analyzed the data; drafted the manuscript for intellectual content |
| Junjie Wu, PhD | Department of Radiology, Emory University. Atlanta, GA | Author | Major role in data acquisition and analysis |
| Deqiang Qiu | Department of Radiology, Emory University. Atlanta, GA | Author | Designed the study; major role in imaging data acquisition and analysis |
| Jennifer Howell | Department of Neurology, Emory University. Atlanta, GA | Author | Major role in data acquisition and analysis |
| Monica W. Parker, MD | Department of Neurology, Emory University. Atlanta, GA | Author | Major role in data acquisition |
| Jessica A Turner, PhD | Psychology Department, Georgia State University. Atlanta, GA | Author | Assisted in the conceptualization of the study; edited the manuscript for critical intellectual content |
| William Hu, MD, PhD | Department of Neurology, Emory University. Atlanta, GA | Author | Major role data acquisition; assisted in the conceptualization of the study; analyzed the data; edited the manuscript for critical intellectual content |

